# Unbiased antimicrobial resistance detection from clinical bacterial isolates using proteomics

**DOI:** 10.1101/2020.11.17.386540

**Authors:** Christian Blumenscheit, Yvonne Pfeifer, Guido Werner, Charlyn John, Andy Schneider, Peter Lasch, Joerg Doellinger

**Author notes:** Correspondence: Joerg Doellinger).

## Abstract

Antimicrobial resistance (AMR) poses an increasing challenge for therapy and clinical management of bacterial infections. Currently, antimicrobial resistance detection often relies on phenotypic assays, which are performed independently from species identification. Although genomics-based approaches are increasingly being proposed as possible alternatives for resistance detection, the analysis of proteins should be superior to gene or transcript sequencing when it comes to phenotype prediction from molecular data as the actual resistance against antibiotics is almost exclusively mediated by proteins. In this study, we present a unbiased proteomics workflow for detecting both, bacterial species and AMR related proteins in the absence of secondary antibiotic cultivation in less than 4 h from a primary culture. The method was validated using a sample cohort of 7 bacterial species and 11 AMR determinants represented by 13 protein isoforms which resulted in a sensitivity of 98 % (100 % with vancomycin inference) and a specificity of 100 % with respect to AMR determinants. This proof-of concept study suggests a high application potential of untargeted proteomics in clinical microbiology.

## Introduction

The rapid emergence and spread of antimicrobial resistant bacteria is a major threat for global health (WHO, 2014). Therefore, the rational use of antibiotics is of utmost importance, which in turn depends on fast and accurate detection of clinically relevant resistances (O’Neill, 2016; WHO, 2015). The most important results in guiding antimicrobial therapy are a correct species identification and a reliable antimicrobial susceptibility testing (AST). Currently no method is applied for routine diagnostics of patient samples, which addresses both needs simultaneously. The last decade has witnessed a revolution in routine clinical microbial diagnostics. MALDI-ToF MS enables today rapid, accurate and cost-effective biotyping at high sample throughput (Patel, 2015; van Veen *et al*, 2010). On the contrary, the determination of antibiotic resistance relies mainly on the analysis of bacterial growth in the presence of antibiotics. This phenotypic approach is often complemented by molecular analysis of targeted genes using polymerase chain reaction (PCR). Due to the high relevance of antimicrobial resistance detection, many different methods are currently under development with the aim at reducing the time-to-results and improving the accuracy (van Belkum *et* al, 2020; van Belkum & Dunne, 2013). These methods split into two main categories. The first approach aims to improve phenotypic methods by increasing the sensitivity of detecting bacterial growth at different conditions and so reduce time to results (Dubourg *et al*, 2018). The second approach uses molecular analysis methods to predict antibiotic resistance from molecular signatures and is currently dominated by next generation sequencing (NGS) (Su *et al*, 2019). Molecular analysis using ‘omics’-technologies offers the potential for accurate prediction of every known resistance phenotype without the need for parallel cultivation in the presence of different antibiotics and without its inherent limitations in transferring the results from the specialized lab cultures into the clinics. In theory, the analysis of proteins should be superior to gene or transcript sequencing when it comes to phenotype prediction as resistances are almost exclusively mediated by proteins (Alcock *et al*, 2020). Therefore, mass spectrometry-based proteomics is a promising candidate as a “one-for-all” solution for addressing the above mentioned needs in clinical microbiology (Welker & van Belkum, 2019). However, several limitations have hampered the breakthrough of this technology so far. Direct detection of the actual proteins conferring resistance by whole cell mass spectrometry biotyping using MALDI-ToF MS is hard to imagine even in the further future (Welker & van Belkum, 2019). Therefore MALDI-ToF MS is currently evaluated as a readout technique for phenotypic antimicrobial susceptibility testing (AST) (Correa-Martínez *et al*, 2019; Lange *et al*, 2014; Nix *et al*, 2020; Sparbier *et al*, 2016). In contrast, liquid chromatography coupled to tandem-mass spectrometry (LC-MS/MS) offers the potential for molecular detection of all known antimicrobial resistance determinants in combination with bacterial species identification. However, no such unbiased LC-MS/MS workflow exists so far. Instead, the majority of AMR-related proteomics studies rely on targeted analysis of selected peptides representing AMR determinants (Cecchini *et al*, 2018; Charretier *et al*, 2015; Foudraine *et al*, 2019; Hassing *et al*, 2016; Strich *et al*, 2019). The results of this studies are quite promising, as they prove that sensitive and specific detection at least of some AMR phenotypes based on proteins is possible. Untargeted proteomics on the other hand has not been evaluated systematically for unbiased antimicrobial resistance prediction most probably due to major technical limitations (Chen *et al*, 2019).

In this study, we present an untargeted LC-MS/MS-based proteomics workflow, which was developed to address the special requirements of antimicrobial resistance detection in clinical microbiology. The proposed workflow not only predicts AMR phenotypes from clinical isolates in an unbiased manner with high sensitivity and specificity using large-scale sequence databases but allows also to simultaneously identify the bacterial species.

## Results

### General workflow

In this study, a proteomic workflow is presented, which enables bacterial species identification and antimicrobial resistance detection (See Figure 1 for an illustration) simultaneously. The method involves preparation of bacterial isolates cultivated in the absence of antibiotics using the filter-aided SPEED protocol (fa-SPEED)(Doellinger *et al*, 2020b). This method is rapid (∼ 30 min), convenient, automatable, and even more important universal for all types of microorganisms, including spore-forming Gram-positive bacteria. The LC-MS/MS measurement relies on data-independent acquisition (DIA), which circumvents the protein detection bias caused by the stochastic nature of data-dependent acquisition (DDA). This strategy allows collection of proteomes with deep coverage (Doellinger *et al*, 2020a). However, it introduces big challenges for data analysis, which need to be covered by tailor-made concepts. In our workflow the peptide identification is based on *in silico* spectral library prediction and neural network supported data analysis using DIA-NN (Demichev *et al*, 2019). However, this approach cannot be used to identify bacteria from very large sequence databases, such as all bacterial entries of the UniprotKnowledgebase (O. n. s. o. r. t. i. u. m. UniProt, 2019). Therefore, we utilized a LC-MS^1^ based method for microbial species identification, which has been adopted from MALDI-ToF MS biotyping and relies on LC-MS^1^ pattern analysis (Lasch *et al*, 2020). The proteome of the identified species is then employed for predicting a sample-specific spectral library, which are in turn used as a background for peptide identification from the Comprehensive Antibiotic Resistance Database (CARD) (Alcock *et al*., 2020). The results are interpreted by means of an in-house R script, named rawDIAtect, which reports the AMR phenotype. The performance of the proposed workflow was evaluated using a panel of 29 bacterial isolates covering 7 species and 11 different AMR determinants represented by 13 protein isoforms, including multidrug-resistant clinical isolates and trans-conjugated bacteria (Table 1).

**Table 1:**
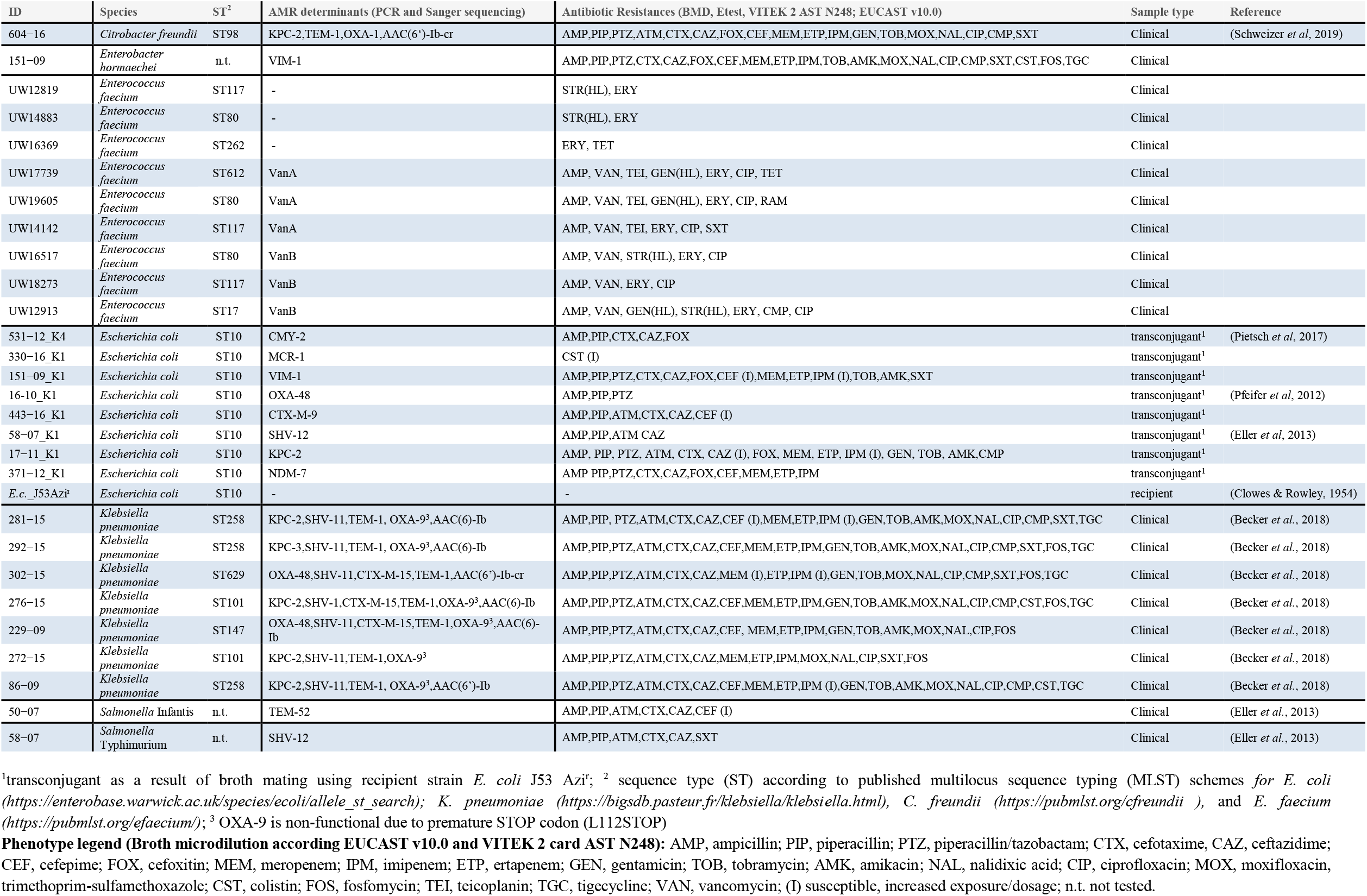
List of bacterial isolates (clinical isolates and transconjugants) with metadata

**Figure 1:**
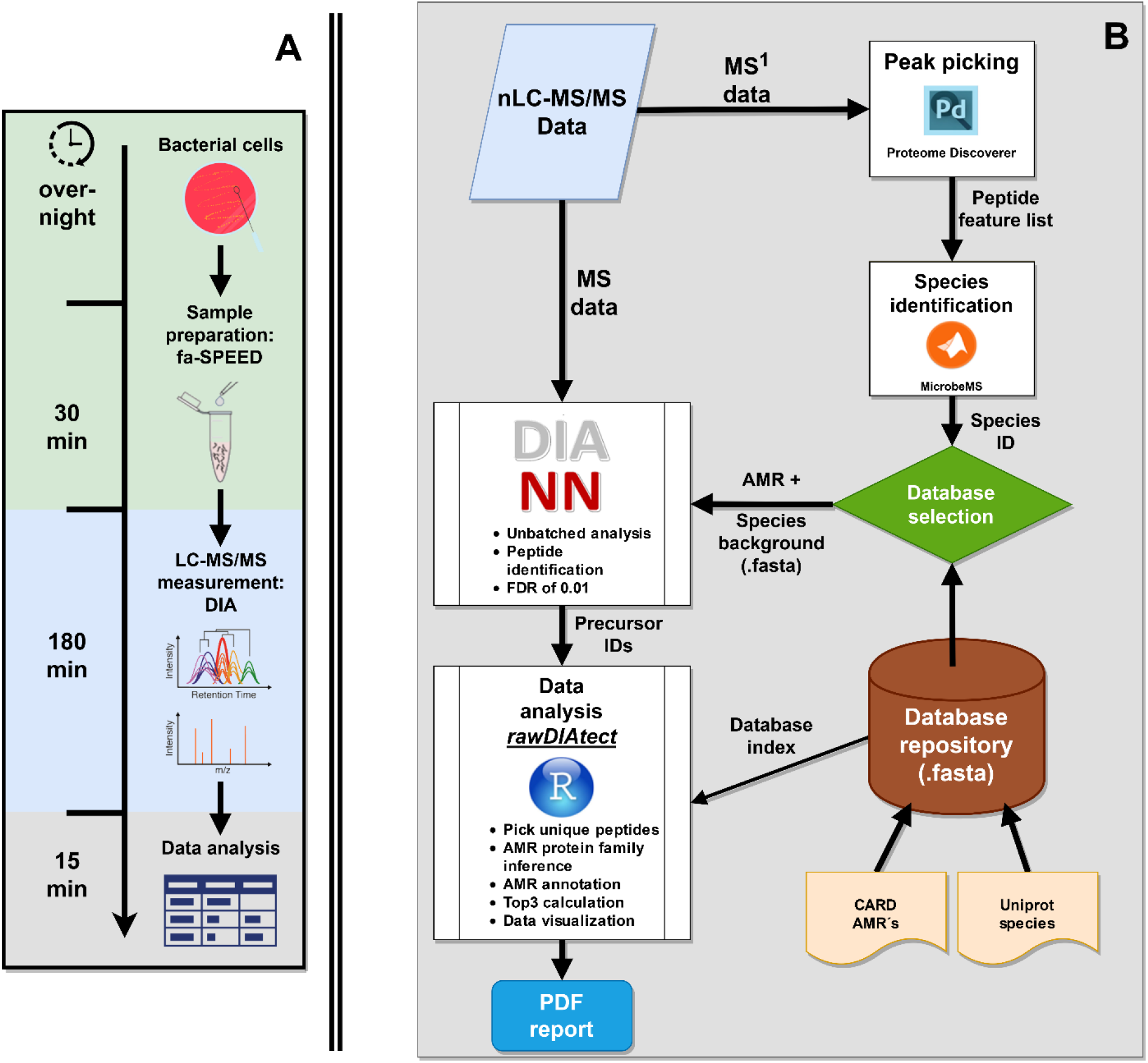
Proteomic workflow of species identification and antimicrobial resistance detection. **A:** Timeline of the whole workflow with cultivation overnight and sample preparation by the fa-SPEED protocol (green). Peptides were measured with a nanoLC setup coupled to a Orbitrap MS system in a data independent manner (blue) followed by the data analysis (gray). **B:** Overview of the data analysis workflow. The MS^1^ data is used for peak picking in the Proteome Discoverer™ using the Minora algorithm. The resulting feature list is then used for species identification with MicrobeMS. The species ID is used to select a species background database from UniProt. This species background database is combined with the CARD AMR database. The combined fasta was put into *DIA-NN* to analyze the MS^2^ data resulting in a list of found peptides. This peptide hit list is then analyzed by the R script *rawDIAtect* to detect AMR’s which give a pdf report of all found resistances.

### Species identification based on LC-MS^1^ biotyping

The LC-MS biotyping approach enables simple and rapid bacterial identification at the species level from very large sequence databases. The method relies on MS^1^ data and thus can be simply integrated into the analysis workflow of DIA data. The results of the LC-MS^1^ biotyping are summarized in Table 2.

**Table 2:**
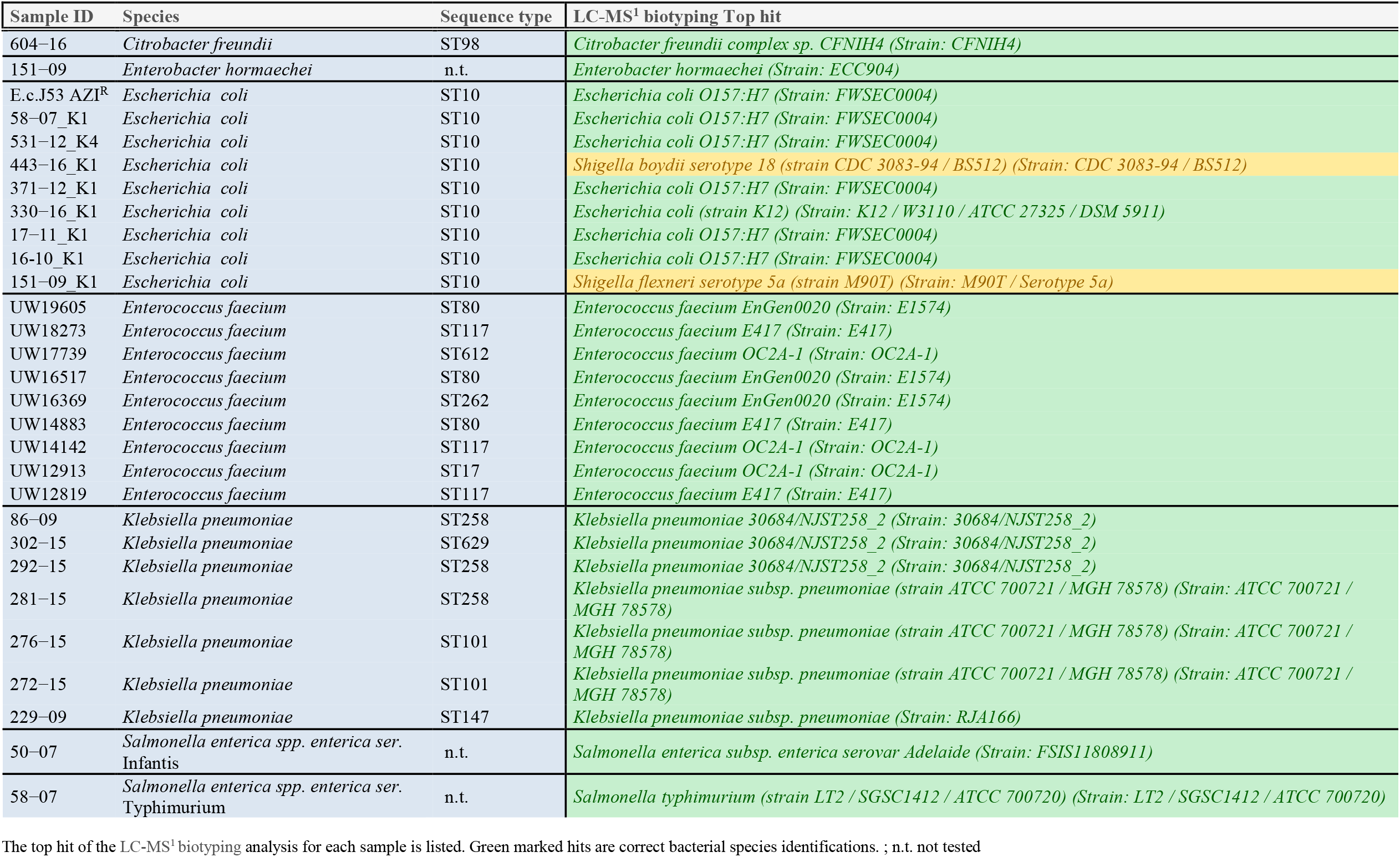
Results of MicrobeMS

| Sample ID | Species | Sequence type | LC-MS<sup>1</sup> biotyping Top hit |
| --- | --- | --- | --- |
| 604-16 | *Citrobacter freundii* | ST98 | *Citrobacter freundii* complex sp. CFNIH4 (Strain: CFNIH4) |
| 151-09 | *Enterobacter hormaechei* | n.t. | *Enterobacter hormaechei* (Strain: ECC904) |
| E.c.J53 AZI<sup>R</sup> | *Escherichia coli* | ST10 | *Escherichia coli* O157:H7 (Strain: FWSEC0004) |
| 58-07_K1 | *Escherichia coli* | ST10 | *Escherichia coli* O157:H7 (Strain: FWSEC0004) |
| 531-12_K4 | *Escherichia coli* | ST10 | *Escherichia coli* O157:H7 (Strain: FWSEC0004) |
| 443-16_K1 | *Escherichia coli* | ST10 | *Shigella boydii* serotype 18 (strain CDC 3083-94 / BS512) (Strain: CDC 3083-94 / BS512) |
| 371-12_K1 | *Escherichia coli* | ST10 | *Escherichia coli* O157:H7 (Strain: FWSEC0004) |
| 330-16_K1 | *Escherichia coli* | ST10 | *Escherichia coli* (strain K12) (Strain: K12 / W3110 / ATCC 27325 / DSM 5911) |
| 17-11_K1 | *Escherichia coli* | ST10 | *Escherichia coli* O157:H7 (Strain: FWSEC0004) |
| 16-10_K1 | *Escherichia coli* | ST10 | *Escherichia coli* O157:H7 (Strain: FWSEC0004) |
| 151-09_K1 | *Escherichia coli* | ST10 | *Shigella flexneri* serotype 5a (strain M90T) (Strain: M90T / Serotype 5a) |
| UW19605 | *Enterococcus faecium* | ST80 | *Enterococcus faecium* EnGen0020 (Strain: E1574) |
| UW18273 | *Enterococcus faecium* | ST117 | *Enterococcus faecium* E417 (Strain: E417) |
| UW17739 | *Enterococcus faecium* | ST612 | *Enterococcus faecium* OC2A-1 (Strain: OC2A-1) |
| UW16517 | *Enterococcus faecium* | ST80 | *Enterococcus faecium* EnGen0020 (Strain: E1574) |
| UW16369 | *Enterococcus faecium* | ST262 | *Enterococcus faecium* EnGen0020 (Strain: E1574) |
| UW14883 | *Enterococcus faecium* | ST80 | *Enterococcus faecium* E417 (Strain: E417) |
| UW14142 | *Enterococcus faecium* | ST117 | *Enterococcus faecium* OC2A-1 (Strain: OC2A-1) |
| UW12913 | *Enterococcus faecium* | ST17 | *Enterococcus faecium* OC2A-1 (Strain: OC2A-1) |
| UW12819 | *Enterococcus faecium* | ST117 | *Enterococcus faecium* E417 (Strain: E417) |
| 86-09 | *Klebsiella pneumoniae* | ST258 | *Klebsiella pneumoniae* 30684/NJST258_2 (Strain: 30684/NJST258_2) |
| 302-15 | *Klebsiella pneumoniae* | ST629 | *Klebsiella pneumoniae* 30684/NJST258_2 (Strain: 30684/NJST258_2) |
| 292-15 | *Klebsiella pneumoniae* | ST258 | *Klebsiella pneumoniae* 30684/NJST258_2 (Strain: 30684/NJST258_2) |
| 281-15 | *Klebsiella pneumoniae* | ST258 | *Klebsiella pneumoniae* subsp. *pneumoniae* (strain ATCC 700721 / MGH 78578) (Strain: ATCC 700721 / MGH 78578) |
| 276-15 | *Klebsiella pneumoniae* | ST101 | *Klebsiella pneumoniae* subsp. *pneumoniae* (strain ATCC 700721 / MGH 78578) (Strain: ATCC 700721 / MGH 78578) |
| 272-15 | *Klebsiella pneumoniae* | ST101 | *Klebsiella pneumoniae* subsp. *pneumoniae* (strain ATCC 700721 / MGH 78578) (Strain: ATCC 700721 / MGH 78578) |
| 229-09 | *Klebsiella pneumoniae* | ST147 | *Klebsiella pneumoniae* subsp. *pneumoniae* (Strain: RJA166) |
| 50-07 | *Salmonella enterica* spp. *enterica* ser. *Infantis* | n.t. | *Salmonella enterica* subsp. *enterica* serovar *Adelaide* (Strain: FSIS11808911) |
| 58-07 | *Salmonella enterica* spp. *enterica* ser. *Typhimurium* | n.t. | *Salmonella typhimurium* (strain LT2 / SGSC1412 / ATCC 700720) (Strain: LT2 / SGSC1412 / ATCC 700720) |

The top hit of the LC-MS<sup>1</sup> biotyping analysis for each sample is listed. Green marked hits are correct bacterial species identifications. ; n.t. not tested

Out of the 29 samples tested 27 were correctly identified at the species level. Only two *E. coli* samples (transconjugants 443-16_K1 and 371-12_K1) were misclassified as *Shigella boydii* and *Shigella flexneri*, respectively. *Shigella* species are closely related to *Escherichia coli* and some methods, including MALDI-ToF MS, cannot be used for reliable differentiation of both species (Khot & Fisher, 2013; Paauw *et al*, 2015), which is a well-known issue in microbiology. Furthermore, the serovar of the *Salmonella enterica* isolate 50-07 was misclassified. The use of peptide mass profiles from LC-MS^1^ data for species identification is a quite new concept, which has an identification rate of 93 % within this dataset. This is well within the range of reported values for MALDI-ToF biotyping, which is a matured and already legally approved technique (Hou *et al*, 2019). Presumably, further improvements of the data analysis workflows for LC-MS^1^-based biotyping should enable even higher identification rates in the future, e.g. considering the fact, that no well-curated database for this purpose is currently available.

### Sensitivity of AMR detection by proteomics

AMR detection by proteomics relies on the assumption that the resistance determinants are expressed even in the absence of antibiotics. We hypothesized, that the detection of proteins conferring resistance is mainly a matter of sensitivity. The proposed experimental workflow provides sufficient analytical sensitivity to cover nearly complete proteomes of clinical isolates. This hypothesis was tested for a panel of 30 samples (29 isolates in 30 measurements, double *E. coli* negative control), which were characterized previously using phenotypic AST, PCR and Sanger sequencing (Table 1).

AMR detection by proteomics was based on the unmodified whole CARD (Full CARD, see above) without filtering short sequences or redundancies. Figure 2 compares outcomes of LC-MS/MS-based AMR-related protein family detection (y-axis) with the corresponding PCR/Sanger reference data (x-axis). The correctness of MS identifications is color coded as either verified by PCR (green), additional (dark orange) or missing (purple). The number of unique peptides of each AMR protein family is denoted in each tile. In total, 43 out of 45 expected proteins conferring resistance were detected by MS. Only VanB ligase was missing in one *E. faecium* samples (UW16517). However, in this sample vancomycin resistance was confirmed by detection of both parts of the two-component system VanS_B_ and VanR_B_ (Figure 2). Furthermore, drug classes associated with protein isoforms were compared with the phenotypic data (Table S 1). The resulting sensitivity was 96 % considering samples, 91 % considering drug classes and 98 % with respect to AMR protein families using a strict matching to genes identified by PCR. When the presence of the VanB protein in the *E. faecium* samples UW16517 is inferred from the expression of its regulatory system VanS_B_/VanR_B_, the sensitivity is 100 % at all levels (Figure 3). We also observed 21 additional protein hits, which were not detected by PCR/sanger sequencing (Figure 2). Among the additional hits, AAC(6)-Ii was found in all *E. faecium* isolates. The respective resistance gene is known to be intrinsic in this species and was therefore not tested by PCR. (Galimand *et al*, 2011; Hollenbeck & Rice, 2012). Due to the nature of the PCR detection being a targeted method, which is not able to cover the heterogeneity of all AMR proteins, this discrepancy in comparison to the proteomics results is expectable. The results of the AMR-related protein identifications including their isoform information are summarized in supplementary Table S 2.

**Figure 2:**
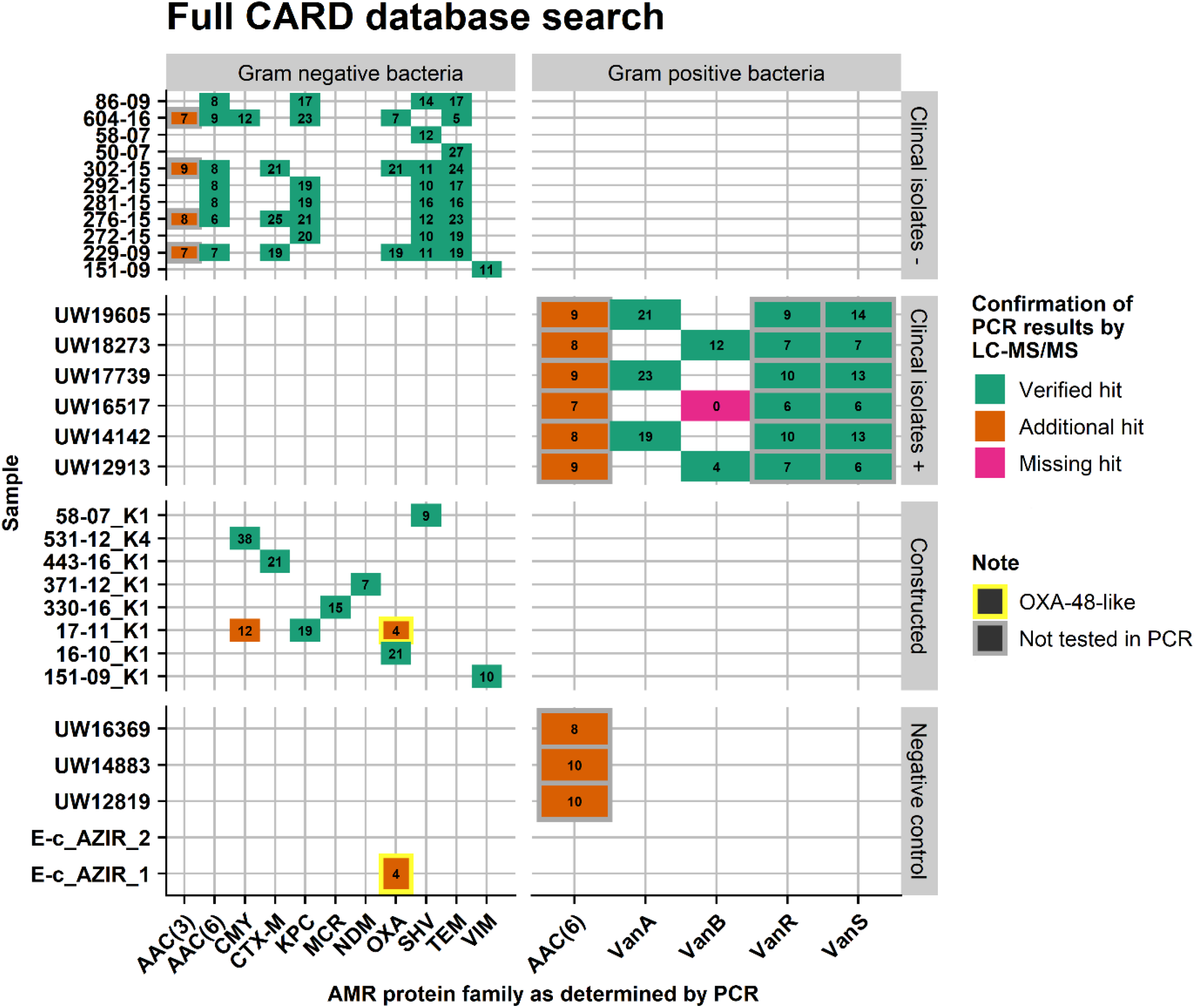
Comparison of AMR protein family detected by proteomics using Full CARD search, and results of PCR/sanger sequencing as the reference method. In green are plotted all proteomic hits verified by PCR. Additional hits of proteomics are plotted in dark orange and missing hits are plotted in pink. The numbers denote the count of unique peptides detected by MS. One missing hit was the Van ligase B, however proteomics was able to detect the corresponding components of the regulatory system VanB_R_ and VanB_S_, which were not analyzed by PCR. This two-component system was detected in all VanB and VanA positive *E. faecium* samples and confirms the presence of a van ligase by the molecular function of the van gene cluster (see text for details).

**Figure 3:**
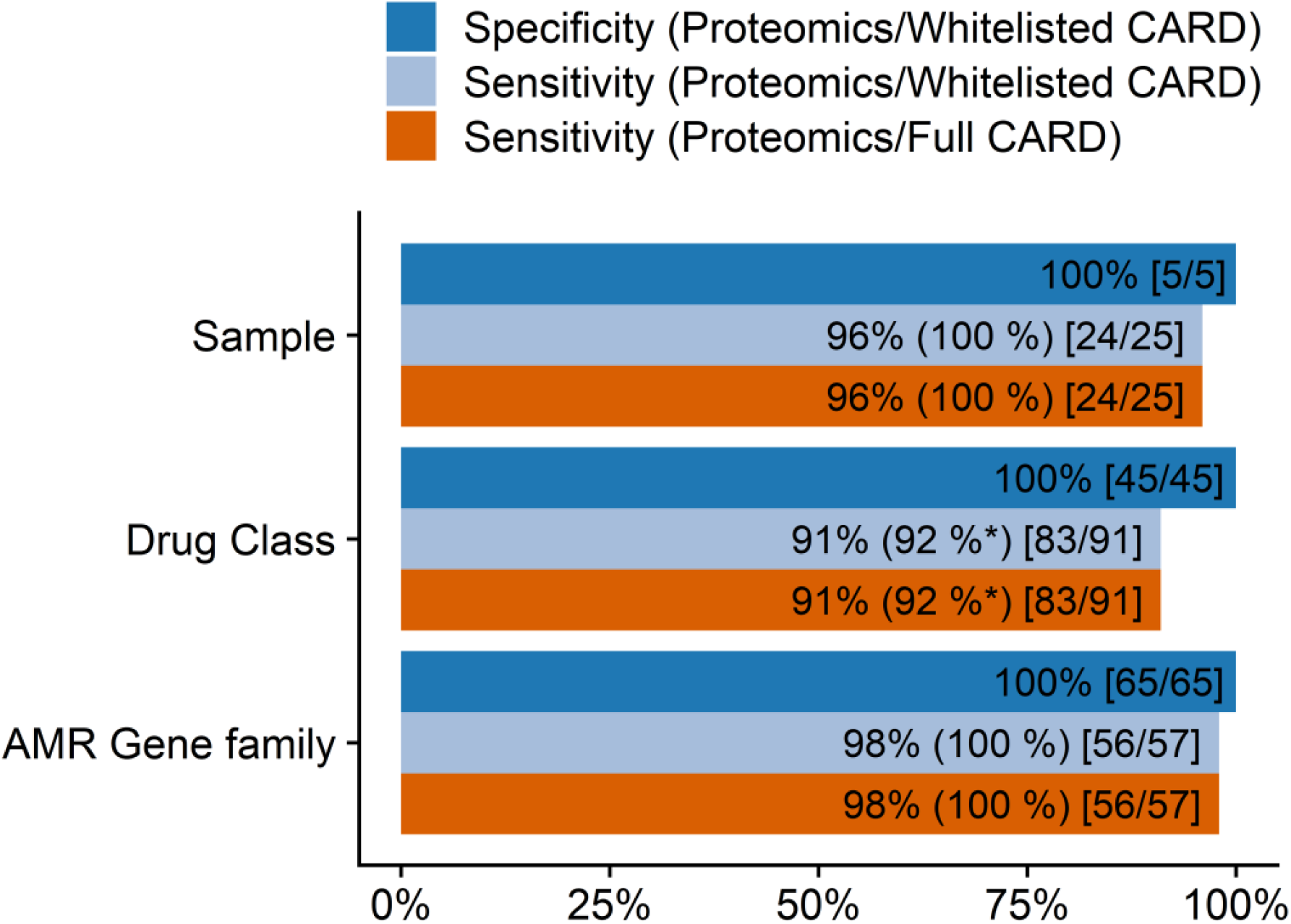
Sensitivity / specificity of AMR detection by proteomics at the sample, drug class and AMR gene level. **Sensitivity** was calculated by the formula TP/(TP + FN) using the results of the Full and Whitelisted CARD in comparison to PCR/Sanger sequencing. **Specificity** was calculated according to TN / (TN + FP) using the results of the Whitelisted CARD search in comparison to PCR/sanger sequencing and NGS tools Abricate (https://github.com/tseemann/abricate, v1.0.1) and Kleborate (https://github.com/katholt/Kleborate, v1.0.0). The sensitivity value in the bracket is calculated after inference of vancomycin resistance by the presence of the van ligase regulatory system (see results and discussion for details). TP/FP - true, or false positive assignment, respectively. TN/FN - true, or false negative assignment. **Definition of TP and TN at the sample level:** A sample was considered as TP only in cases where all AMR protein families (as determined by PCR/Sanger sequencing) were correctly assigned (FN otherwise). Vice versa, a sample was defined as TN at the sample level if no AMR family was determined in the control samples (FP otherwise). One false negative (FN) assignments was observed (missed VanB detection, see text for details). **TP and TN at the drug class level:** A found drug class, was counted as TP, if matched to a phenotype of the sample. 91 drug classes hits over all measurements were observed. One FN, the missing vancomycin hit was observed and 7 predictions can only be potently matched due to combination of multiple resistances. A TN was only counted if no drug class was found. In total 9 drug classes could be found per negative measurement. * Prediction of Drug class considering all possible protein isoform hits including vancomycin. **TP / TN at the gene family level:** A TP was counted for every correct AMR gene family hit given by PCR and missing hits as FN. One missing VanB family hit was observed. In total 57 AMR gene family hits could be found over all measurements. TN was only counted if no hit could be found for any possible AMR determinant (13) per negative measurement.

### Specificity of AMR detection by proteomics

The specificity of the proteomic approach was analyzed after reduction of the CARD to contain only the potential PCR targets (Figure 4, Whitelisted CARD approach). In contrast to Figure 2, we compared in the example of Figure 4 the MS hits not only against the PCR results, but also to NGS data (Table S 2) in order to compensate for the limitations of the PCR. The results of 30 LC-MS measurements on the y-axis are matched against the possible PCR and NGS hits on the x-axis Again, 43 out of 45 expected proteins conferring resistance were identified, while VanB ligase was again missing in one *E. faecium* samples. The AAC(6)-Ib-like hits in all three negative controls of our *E. feacium* samples in the Full CARD approach were not detected anymore by the Whitelisted approach. Furthermore, no AMR determinants were detected in the *E. coli* negative controls. AAC(3)-like proteins were not tested in the Whitelisted CARD approach. The additional hits in both isolates 17/11 K1 and 330-16-K1 identified by the Full CARD approach were also not detected anymore, indicating good agreement between PCR and LC-MS results if equal reference sequences are used. In total, no resistance-related proteins were identified in the 5 negative control samples, which resulted in a specificity of 100 % considering samples, drug classes and AMR protein families (Figure 3).

**Figure 4:**
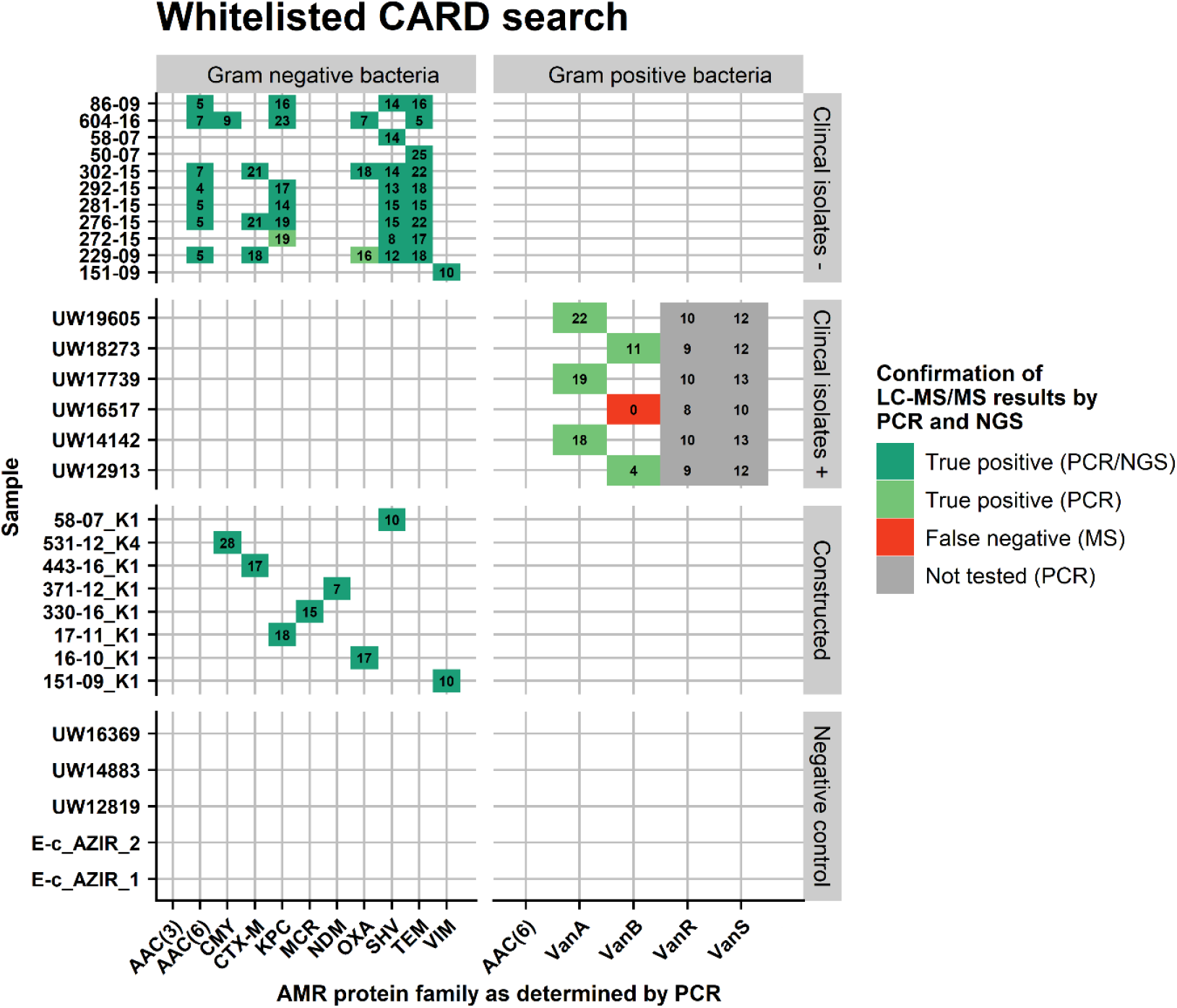
Comparison of AMR protein family detected by proteomics using Whitelisted CARD search approved by PCR/sanger sequencing and NGS as the reference method. Proteomic hits verified by PCR and NGS are plotted in dark green. Additional hits of proteomics, which were verified only by PCR are plotted in light green and missing hits are plotted in red. The numbers stand for the count of unique peptides detected by MS. One missing hit was the Van ligase B, however proteomics was able to detect the corresponding components of the regulatory system VanR and VanS, which were not analyzed by PCR (Gray). This two-component system was detected in all VanB and VanA positive *E. faecium* samples and confirms the presence of a van ligase by the molecular function of the van gene cluster. No additional hits of proteomics, which could not be confirmed by either PCR or NGS (Abricate, https://github.com/tseemann/abricate, v1.0.1 and Kleborate, https://github.com/katholt/Kleborate, v1.0.0) were detected.

## Discussion

Accurate identification of the microbial species and correct determination of antimicrobial resistance pattern are of paramount importance to assure adequate treatment of patients suffering from bacterial infections. The aim of this proof-of-concept study was thus to establish an untargeted proteomics workflow suitable for the comprehensive characterization of pathogenic bacteria that includes both, species identification and AMR type determination. In contrast to culture-based phenotypic methods, this molecular proteomic-based analysis of clinical isolates has the potential for accurate prediction of every known resistance phenotype in a truly untargeted manner. Currently, genomics is the only ‘omics’-technology which has been systematically evaluated for this purpose, although the analysis of proteins could offer some advantages due to its close relationship to the phenotype. Proteins directly mediate almost all resistant phenotypes and can be therefore considered ideal targets for antimicrobial susceptibility testing (AST). The potential of proteomics for AMR detection in general has been shown earlier using targeted analysis of distinct proteins. However, as antibiotic resistance databases, such as the Comprehensive Antibiotic Resistance Database (CARD), which currently includes > 1600 AMR protein families, are continuously growing and many of these gene families are highly heterogeneous a targeted approach for comprehensive AMR detection seems not effective (Alcock *et al*., 2020). So far major technological challenges have prevented untargeted proteomics from being a competitive alternative to NGS. In this study, we propose a proteomics workflow, which combines several recent technological developments into a workflow strategy, which was not possible until recently. The strategy is based on the *in-silico* prediction of sample-specific peptide spectral libraries as well as universal resistance-related peptide spectral libraries obtained by deep learning. This enables the use of DIA-based proteomics, which records the entity of peptide fragments in a sample above its sensitivity threshold in an unbiased manner with deep proteome coverage. The prediction of sample-specific background libraries for AMR detection is based on the results of a recently introduced strategy for biotyping bacteria using LC-MS^1^ data, whose recording can easily be integrated into DIA measurements (Lasch *et al*., 2020). This approach resulted in an identification accuracy of 93 % with a runtime of ∼ 1 min on a decent desktop PC using a database with 8540 bacterial entries. The workflow further involves very rapid (∼30 min) and universal sample preparation using acid extraction, without the need for protocol adaptation to certain bacterial species. Potentially, this method, termed SPEED, should enable to fully automate sample preparation including bacterial cell lysis in the future, as it uniquely omits the need for physical disruption (Doellinger *et al*., 2020a).

The performance of this workflow for AMR detection was tested with a panel consisting of 20 clinical isolates and 9 *E. coli* transconjugants containing a broad range of beta-lactamases (SHV-1/-11/-12, TEM-1/-52, CMY-2, CTX-M-15/-9, KPC-2/-3, NDM-7, OXA-48/-1, VIM-1), the aminoglycoside acetyltransferases AAC(6)-like, *vanA*- and *vanB*-type vancomycin resistances and plasmid mediated colistin resistance (Mcr). The sensitivity was found to be 98 % (100% with vancomycin inference) and the specificity was 100%, with respect to AMR-related protein families (cf. Figure 3). This demonstrates that antimicrobial resistance can be sensitively detected by protein analysis even in the absence of antibiotics during cultivation and demonstrates the high potential of untargeted proteomics for routine diagnostics in clinical microbiology.

The current workflow used in this proof-of-concept study takes ∼ 4 h from bacteria to results, of which ∼ 3.3 h is needed for the actual LC-MS/MS measurement. Recent advancements in MS instrumentation in combination with liquid chromatography using higher flow rates have dramatically increased the proteome coverage when using short LC gradient lengths (< 30 min) (Bache *et al*, 2018; Meier *et al*, 2018; Messner *et al*, 2020). Therefore, we expect, that the LC-MS/MS measurement time needed to provide sufficient depth for clinical microbiology should massively decrease in the future and thus increase sample throughput. Ultimately, this could end up in the unbiased characterization of bacteria directly from clinical specimen without any cultivation.

An important consideration for routine clinical applications of this workflow is the establishment of curated and validated peptide databases as the use of unprocessed publicly available protein sequence databases suffers from entries with low quality. Such peptide databases could also include information on discriminatory sequences, which could be used to distinguish clinically relevant protein isoform families, such as ESBL and non-ESBL variants of SHV and TEM, which is not possible with the current workflow. Antimicrobial resistance is often associated with a high genetic variability. However, the clinical relevance of the vast majority of genetic variants is unknown. RawDIAtect was able to discriminate between AMR protein subfamilies like carbapenemase producing OXA 48-like proteins and non-carbapenemase producing OXA 1-like proteins and further reports all possible isoforms for the detected peptide sequences as sequence coverage for most proteins is incomplete in many instances.

The output of rawDIAtect already contains information on the protein abundance. This data could potentially be used for the categorization of isolates into susceptible, intermediate or resistant strains just as the current use of minimum inhibitory concentration (MIC) breakpoints and so enable antimicrobial susceptibility testing (AST). A correlation between abundance of resistance determinants and MICs has already been demonstrated before for RNA & NGS data (Bhattacharyya *et al*, 2019; Khaledi *et al*, 2016; Nguyen *et al*, 2019). These further improvements of the workflow could potentially turn this proof-of concept workflow into a method ready for routine applications in clinical microbiology.

## Materials and Methods

### Bacterial Cultivation

A panel of antimicrobial resistant bacteria consisting of 29 isolates with 7 species including Gram-negative (*Escherichia coli* (n=9), *Klebsiella pneumoniae* (n=7), *Citrobacter freundii* (n=1), *Enterobacter hormaechei* (n=1), *Salmonella enterica ser*. Infantis (n=1), *Salmonella enterica ser*. Typhimurium (n=1)) and Gram-positive bacteria (*Enterococcus faecium* (n=9)) was analyzed in 30 LC-MS/MS runs (double *E. coli* negative control) (Table 1). The Gram-negative isolates contained various β-lactamases including carbapenemases and extended-spectrum β-lactamases (ESBL). Some isolates additionally carried aminoglycoside acetyltransferases (AAC(6)-Ib-like) and plasmid-mediated colistin resistance (Mcr-1). The *E. coli* isolates were β-lactamase producing transconjugants that were obtained from clinical donor strains by mating experiments using the sodium azide-resistant recipient strain *E. coli* J53 Azi^r^. The enterococci contained various vancomycin resistance gene clusters (*vanA*-type, *vanB*-type) including genes encoding a VanA or a VanB ligase, respectively. Cells were streaked under sterile conditions on Mueller Hinton II Agar (Becton Dickinson, Heidelberg, Germany) and aerobically grown at 37°C overnight for at least 15 h prior to proteomic sample preparation.

### Antimicrobial susceptibility testing (AST)

Antimicrobial susceptibilities of all 29 isolates were determined by the use of broth microdilution (BMD) and VITEK 2 card AST N248 according to DIN58940 and EUCAST guidelines v10.0 (www.eucast.org). In total, 19 antibiotics were tested, including ampicillin, piperacillin, piperacillin/tazobactam, cefotaxime, ceftazidime, cefoxitin, cefepime, meropenem, imipenem, gentamicin, amikacin, tobramycin, nalidixic acid, ciprofloxacin, moxifloxacin, fosfomycin, tigecycline, sulfamethoxazole/trimethoprim and colistin. Additionally, ETEST^®^ with imipenem and ertapenem (bioMérieux, Nuertingen, Germany) were performed for selected isolates. AST of enterococcal isolates was determined with BMD and according to EUCAST standards and guidelines. For isolates that do not have a clinical breakpoint, corresponding ECOFF values were applied to discriminate the wildtype population from non-wildtype (= resistant).

### PCR-based detection of resistance genes

The presence of various β-lactamase genes (*bla*_NDM-like_, *bla*_VIM-like_, *bla*_OXA-48-like_, *bla*_KPC-like_, *bla*_CTX-M-1-2-9group_, *bla*_TEM-like_, *bla*_SHV-like_, *bla*_CMY-like_, *bla*_OXA-1-2-9-10group_,) was tested by PCR and subsequent Sanger sequencing using previously described primers and protocols (Gröbner *et* al, 2009; Pfeifer *et al*, 2011; Turton *et al*, 2006). Additionally, PCR screening and subsequent Sanger sequencing for genes contributing to resistance to aminoglycosides (*aac(6’)-Ib-like*) and colistin (*mcr-1*) was performed as described elsewhere (Liu *et al*, 2016; Park *et al*, 2006). The presence of vancomycin resistance gene clusters was determined by detecting corresponding ligase genes *vanA* and *vanB* by a multiplex PCR according to an accredited in house protocol (Werner *et al*, 2012).

### NGS-based detection of resistance genes

NGS data were analyzed in order to identify resistance genes and thus confirm PCR and LC-MS/MS results. Sequence data were already publicly available for the *K. pneumoniae* isolates (Becker *et al*, 2018). Both *Salmonella* isolates as well as the *C. freundii* and the *E. hormaechei* complex isolates were sequenced in the course of this study with Illumina HiSeq in a paired 150 bp manner with an aimed coverage of 20x by LGC Genomics (Berlin, Germany). The bacterial DNA was extracted using the DNeasy Blood and Tissue kit according to the protocol of the manufacturer (Qiagen, Hilden, Germany). Quantification was carried out using the Qubit dsDNA HS Assay Kit (Invitrogen/ Thermo Fisher Scientific, Karlsruhe, Germany). Library preparation with 1 ng of extracted DNA was done by applying the Nextera XT DNA Library Prep Kit according to the manufacturer’s instructions (Illumina, San Diego, CA). Libraries were subjected to whole-genome sequencing using an Illumina^®^ MiSeq benchtop devise (Illumina, San Diego, CA) in paired-end mode (v3 chemistry; 2x 300 bp). All reads were trimmed and filtered using *fastp* on default settings (https://github.com/OpenGene/fastp, v0.20.1) and assemblies were generated de-novo from trimmed and filtered reads by *Unicycler* using the short only read assembler option (https://github.com/rrwick/Unicycler, v0.4.8) without read correction and normal bridging. Detection of AMR relates genes was done with Abricate (https://github.com/tseemann/abricate, v1.0.1) using the CARD database and Kleborate (https://github.com/katholt/Kleborate, v1.0.0). The NGS results are summarized in Table S 2.

### Sample preparation for proteomics

Bacteria were prepared for mass spectrometry by *filter-aided Sample Preparation by Easy Extraction and Digestion* (fa-SPEED) as described previously (Doellinger *et al*., 2020b). In brief, bacterial cells were harvested using an inoculating loop, washed twice with PBS and pelleted for 5 min at 5,000 x g. Pellets were resuspended in trifluoroacetic acid (TFA) (Optima® LC/MS, Thermo Fisher Scientific, Schwerte, Germany) (sample/TFA 1:4 (v/v) and incubated at 70°C for 3 min. Samples were neutralized with 2M TrisBase using 10 x volume of TFA and further incubated at 95°C for 5 min after adding Tris(2-carboxyethyl)phosphine (TCEP) to a final concentration of 10 mM and 2-chloroacetamide (CAA) to a final concentration of 40 mM. Protein concentrations were determined by turbidity measurements at 360 nm (1 AU = 0.67 µg/µL) using the NanoPhotometer® NP80 (Implen, Westlake Village, California). Note, that the turbidity values are instrument dependent, for further details see Doellinger *et. al* 2020 (Doellinger *et al*., 2020b). 40 µg proteins were mixed with acetone (Uvasol® for spectroscopy, Merck, Darmstadt, Germany) to obtain a final concentration of 80 % acetone (v/v) and pipetted on an Ultrafree-MC PTFE membrane 0.2 µm filter (Merck Millipore, Burlington, MA). The filter was incubated for 2 min at room temperature and was then centrifuged at 5000 x g for 1 min. The samples were washed successively with 200 µL 80% acetone, 180 µL acetone and 180 µL n-pentane for 1 min at 5000 x g each. The filter was then dried for 2 min under a fume hood to remove all remaining solvents. Afterwards 40 µL digestion buffer (1:10 diluted Rapid Digest buffer (Promega, Fitchburg, WI) in water) containing Rapid Digestion Trypsin (Promega) using a protein/enzyme (w/w) ratio of 10:1 was added. The filter was incubated for 15 min in a thermomixer (Thermo Fisher Scientific, Bremen, Germany) at 70°C covered by a beaker filled with 70 °C hot water. After digestion samples were spun down for 1 min at 5000 x g. The eluted peptide solution was acidified with 10 µL 10% TFA to a pH of ∼2. Concentration was measured with the NanoPhotometer® NP80 (Implen, Westlake Village, CA). All samples were diluted with 0.1 % TFA to obtain a final peptide concentration of 0.25 µg/µL and stored at −80°C.

### Liquid chromatography and mass spectrometry

Peptides were analyzed on an EASY-nanoLC 1200 (Thermo Fisher Scientific, Bremen, Germany) coupled online to a Q Exactive™ Plus mass spectrometer (Thermo Fisher Scientific, Bremen, Germany). 1.25 µg peptides were loaded on a µPAC™ trap column (5 μm pillar diameter, 18 µm pillar length, 100*2 mm bed channel size, 100-200 Å C18, PharmaFluidics, Ghent, Belgium) and then separated on a µPAC™ analytical column (5 μm pillar diameter, 18 µm pillar length, 315 µm*200 cm bed channel size, 100-200 Å C18, PharmaFluidics, Ghent, Belgium) using a stepped 160 min gradient of 80 % acetonitrile (solvent B, Optima LC/MS, Fisher Chemical) in 0.1 % formic acid (solvent A, Optima LC/MS, Fisher Chemical) at 300 nL/min flow rate: 3-10% B in 22 min, 10-33% B in 95 min, 33-49% B in 23 min, 49-80% B in 10 min and 80% B for 10 min. The 200 cm µPAC™ column was kept at 50°C using a butterfly heater (Phoenix S&T, Chester, PA. In order to reduce peptide carry-over, 1 µg *Candida albicans* strain SC5314 digest was loaded on the trap column and subsequently washed using 15 µL wash solution (60 % acetonitrile, 30 % isopropanol and 9.9 % water and 0.1% FA) after each sample. During the equilibration step of the next sample, the wash solution that remained in the trap column was then used to wash the analytical column. The Q Exactive™ Plus was operated in a data-independent (DIA) manner in the m/z range of 350 – 1,150. Full scan spectra were recorded with a resolution of 70,000 using an automatic gain control (AGC) target value of 3 × 10^6^ with a maximum injection time of 100 ms. The Full scans were followed by 69 DIA scans of dynamic window widths using an overlap of 0.5 Th (See window table in supplement for more information). DIA spectra were recorded at a resolution of 17,500 using an AGC target value of 3 × 10^6^ with a maximum injection time of 55 ms and a first fixed mass of 200 Th. Normalized collision energy (NCE) was set to 25 % and default charge state was set to 3. Peptides were ionized using electrospray with a stainless-steel emitter, I.D. 30 µm, (Proxeon, Odense, Denmark) at a spray voltage of 2.1 kV and a heated capillary temperature of 275°C.

### Species identification by LC-MS^1^ data

Identification of microbial species was carried out as outlined by (Lasch *et al*., 2020) with slight modifications. Briefly, peptide (MS^1^) feature lists were extracted from raw DIA data using the Minora algorithm of the *Proteome Discoverer*™ software (v2.2.0388, Thermo-Fisher Scientific). The feature lists were pre-processed and subsequently exported into a format compatible with *MicrobeMS*, a Matlab toolbox (Matlab: The Mathworks, Natick, MA) developed in house for microorganism identification by mass spectrometry (http://wiki.microbe-ms.com). Using MicrobeMS, MS^1^ feature lists were compared against a library of strain-specific *in silico* mass profiles obtained from UniProtKB/Swiss-Prot and UniProtKB/TrEMBL protein sequence data. Ranking lists of correlation, or inter-spectral distance values (i.e. of scores) were obtained that provided information on the taxonomic identity of the organism under study (please see Lasch *et al*, for details). The *in-silico* database (v2.0) consisted of 8580 strain-specific *in silico* profiles, each containing tens of thousands of peptide mass entries. As distance values, variance-scaled Pearson’s product momentum correlation coefficients (Pareto scaling 0.25) were selected, whereby data between 2000 and 4500 Da served as inputs. The results of correlation analysis were scored between 0 (no correlation) and 1000 (identity) and arranged in ranking lists to determine the taxonomic identity of the samples investigated.

### Peptide identification

The reference proteomes with the highest completeness according to their BUSCO values (Waterhouse *et al*, 2018) and with a full genome representation were downloaded from UniProtKB for each of the 7 microbial species. AMR-related proteins in this species-specific databases, were then removed using BLASTp algorithm implemented in *DIAMOND* (v0.9.29, https://github.com/bbuchfink/diamond) (Buchfink *et al*, 2014) with a sequence identity threshold of 95% with respect to any entry of the Comprehensive Antibiotic Resistance Database (CARD v3.0.5, September 2019)(Alcock *et al*., 2020). This modified proteome FASTA is then used as a species-specific background database for detecting AMR-related proteins after merging the databases with all CARD entries. The headers for each protein entry of the CARD, were modified to match our workflow resulting in the following motif: “>CD|Protein name|ARO:XXXXXXX GN=ARO:XXXXXXX”. All LC-MS/MS raw data files were analyzed using the species-specific background databases merged with either the whole unfiltered CARD called “Full CARD” (196.072 entries) or a “Whitelisted CARD” (22 entries), which contains only the protein isoforms, targeted previously by PCR (CMY-2, CTX-M-9/15, KPC-2/3, NDM-1, OXA-1/48, SHV-1/12, TEM-1/52, VIM-1, AAC(6)-lb-cr, VanA-R/S and VanB-R/S). Those protein isoforms were extracted from the Full CARD using Geneious Prime® (v2020.0.5) and consensus sequences were generated by multiple sequence alignment (MSA) using *MAFFT* (v6.240, https://mafft.cbrc.jp/)(Katoh & Standley, 2013; Rozewicki *et al*, 2019).

Peptides in the LC-MS/MS data were identified using DIA-NN (v1.76) (Demichev *et al*., 2019). Samples were analyzed separately for each species in an unbatched manner in conjunction with the unrelated run option. Spectral libraries were predicted using the deep-learning algorithm implemented in DIA-NN with strict trypsin specificity (KR not P) allowing up to one missed cleavage site in the m/z range of 350 – 1,150 for all peptides consisting of 7-30 amino acids of the sample-specific background database and either the “Full” or “Whitelisted” version of CARD. Cysteine residues were assumed to be carbamidomethylated and protein N-terminal methionines were removed. The mass spectra were analyzed using fixed mass tolerances of 10 ppm for MS^1^ and 20 ppm for MS^2^ spectra. Retention time profiling was disabled. The false discovery rate was set to 1 % for precursor identifications and proteins were grouped according to their respective genes.

### Antimicrobial resistance detection and reporting

AMR protein families and isoforms as well as the associated drug classes were detected from the DIA-NN output files using the R (v3.6.0) script *rawDIAtect* developed in house (v0.3, https://github.com/CptChiler/rawDIAtect), which consists of two modules: *rawSplit* and the *“****C****linical* ***R****esistance* ***I****dentification* ***T****ool”* (*CRIT)*. The first module *rawSplit*, was used to split the peptide result file from DIA-NN into single files for each sample. These files were then loaded into the module *CRIT*, which extracted all peptides identified from CARD and annotated these hits with antimicrobial resistance related information such as AMR protein family, drug class and AMR protein subfamily deposited in the ARO index file of CARD Ontology (v3.0.5, September 2019) (Alcock *et al*., 2020). Peptide sequences were then filtered for uniqueness with respect to AMR protein families and for removal of multiple charge states keeping only the charge state with the highest precursor quantity. AMR protein families were inferred from the remaining peptides and were subsequently filtered requiring a minimum of 3 sequences per protein. After this, the abundance of each AMR protein family was calculated from the top 3 precursor intensities (Grossmann *et al*, 2010; Silva *et al*, 2006). The most likely AMR protein isoforms were inferred from ranking all identified isoforms per AMR protein family according to their peptide counts. Protein isoforms of one AMR protein family were matched to their subfamily according to the Beta-lactamase database (BLDB) (Naas *et al*, 2017) and given a new tag “Subfamily” to clear up multiple isoform hits. All findings were collected into a table and visualized with the R package *ggplot2*. All plots were summarized in a single pdf file and the processed peptide tables were separately saved for further investigation (uploaded on Pride).

## Supporting information

Table S 1: Validation of drug classes detected by proteomics with phenotypic resistance detection

Table S 2: AMR protein isoform identifications using different approaches

Table S 3: Isolation windows corresponding to 1.5 data points per peak (160 min gradient)

## Acknowledgments

We thank Sibylle Müller-Bertling, Kirstin Ganske and Kathleen Klaper for excellent technical and microbiological assistance. We also thank Stephan Fuchs for his support writing this paper.

## Author contributions

J.D., A.S., C.B. and P.L. conceived the study and developed the method.

C.B., J.D. and P.L. designed the experiments.

C.B., C.J. and J.D, performed the proteomic experiments.

Analysis of proteomic data was done by C.B., J.D. and P.L.

C.B. developed the R package rawDIAtect.

J.D., YP and GW performed the characterization of the isolates (AST, PCR, Sanger sequencing).

C.B., P.L, J.D., W.G. and Y.P. analyzed the study data and wrote the manuscript.

## Competing interests

J.D., A.S. and P.L. are the inventors of SPEED and have submitted a patent application related to SPEED. The other authors declare that they have no competing interests.

## Data Availability

### Access to proteomics data

The mass spectrometry proteomics data have been deposited to the ProteomeXchange Consortium (http://proteomecentral.proteomexchange.org) via the PRIDE partner repository with the dataset identifiers **PXD022425**. For access to data contact us

### Access to genomic data

All used genomic data have been deposited at the EMBL-EBI database (https://www.ebi.ac.uk/) and can be found under the access ID **ERP123377**.

### Access to code and programs

The R-code of rawDIAtect is available on https://github.com/CptChiler/rawDIAtect. MicrobeMS can be downloaded from https://www.microbe-ms.com/.

#### Abbreviations

AGC: Automatic gain control
AMR: Antimicrobial resistance
ARO: Antibiotic Resistance Ontology
AST: Antimicrobial Susceptibility Testing
BLDB: Beta-lactamase database
BMD: Broth microdilution
CAA: 2-chloroacetamide
CARD: Comprehensive Antibiotic Resistance Database
DB: Database
DDA: Data dependent acquisition
DIA: Data independent acquisition
ESBL: Extended spectrum beta lactamases
ESKAPE: *Enterococcus feacium, Staphylococcus aureus, Klebsiella pneumoniae, Acinetobacter baumannii, Pseudomonas aeruginosa, Enterobacter spp*.
fa-SPEED: filter-aided Sample Preparation by Easy Extraction and Digestion
MALDI-TOF: Matrix-assisted laser desorption/ionization Time-of-flight
MS: Mass spectrometry
MSA: Multiple sequence alignment
NCE: Normalized collision energy
NGS: Next generation sequencing
PBS: Phosphate-Buffered Saline
PCR: Polymerase chain reaction
PRM: Parallel Reaction Monitoring
PTFE: Polytetrafluoroethylene
RKI: Robert Koch Institute
SPEED: Sample Preparation by Easy Extraction and Digestion
TCEP: tris(2-carboxyethyl)phosphine
TFA: Trifluoroacetic acid
WGS: Whole genome sequencing

## Notes

http://proteomecentral.proteomexchange.org/cgi/GetDataset?ID=PXD022425

https://www.ebi.ac.uk/ena/browser/view/PRJEB39816

https://github.com/CptChiler/rawDIAtect

https://www.microbe-ms.com/

